# All-atom protein sequence design based on geometric deep learning

**DOI:** 10.1101/2024.03.18.585651

**Authors:** Jiale Liu, Zheng Guo, Changsheng Zhang, Luhua Lai

## Abstract

The development of advanced deep learning methods has revolutionized computational protein design. Although the success rate of design has been significantly increased, the overall accuracy of *de novo* design remains low. Many computational sequence design approaches are devoted to recover the original sequences for given protein structures by encoding the environment of the central residue without considering atomic details of side chains. This may limit the exploration of new sequences that can fold into the same structure and restrain function design that depends on interaction details. In this study, we proposed a novel deep learning framework, GeoSeqBuilder, to learn the relationship between protein structure and sequence based on rotational and translational invariance by extracting the information from relative locations. We utilized geometric deep learning to fetch the spatial local geometric features from protein backbones and explicitly incorporated three-body interactions to learn the inter-residue coupling information, and then determined the central residue type. Our model recovers over 50% native residue types and simultaneously gives highly accurate prediction of side-chain conformations which gives the atomic interaction details and circumvents the dependence of protein structure prediction tools. We used the likelihood confidence log*P* as scoring function for sequence and structure consistence evaluation which exhibits strong correlation with TM-score, and can be applied to recognize near-native structures from protein decoys pool in protein structure prediction. We have used GeoSeqBuilder to design sequences for two proteins, including thioredoxin and a *de novo* hallucinated protein. All of the 15 sequences experimentally tested can be expressed as soluble monomeric proteins with high thermal stability and correct secondary structures. We further solved one crystal structure for thioredoxin and two for the hallucinated structure and all the experimentally solved structures are in good agreement with the designed models. The two designed sequences for the hallucination structure are novel without any homologous sequences within the latest released database clust30. The ability of GeoSeqBuilder to design new sequences for given protein structures with atomic details makes it applicable, not only for *de novo* sequence design, but also for protein-protein interaction and functional protein design.

## Introduction

Most proteins conduct their functions via folding to specific tertiary structures. Understanding the protein sequence-structure-function relationship provides key information for protein structure prediction and design. Protein structure prediction [1–11] has made awesome development through leveraging the deep learning techniques and the vast protein sequence data. Computational protein *de novo* sequence design, as the reverse folding problem of protein structure prediction, tries to generate novel protein sequences that can fold into the target structure, and creates functions for potential therapeutics, biosensors and biocatalysis applications.

Conventional protein design approaches based on physical energetic functions or statistical potentials [12–14] search for sequences that can fold to the target structure with minimum energy by sampling the protein side-chain rotamers. The rational design process usually involves several cycles of design and experimental testing or even in vitro evolution due to the low success rate of computational design, which is computationally and experimentally demanding. These approaches normally only consider two-body interactions and the generated sequences usually lack divergence. Deep learning, as a kind of high dimensional representative tool, can learn more complex and high order relationships between protein structures and sequences. Computational protein design by using deep learning approaches has attracted much attention in the post AlphaFold2 (AF2) era. Though a number of deep learning based protein sequence design methods [15–26] have been developed, only a few of them have been experimentally tested. Most of the computational design approaches focus on achieving high sequence recovery rate, which result in low diversity of the generated sequences. For example, Anand. et al [22] used a 3D-CNN based method to design protein sequences for TIM-Barrel structure and successfully solved the designed protein structure. The computational efficiency is also relatively low as generating a sequence with 100 residue length usually needs to a few hours. Liu. et al developed ABACUS-R [27] that uses an encoder-decoder framework to update the decoding central residue type with self-consistence. The identity of the designed protein sequences with native structures is about 50%. ProteinMPNN [28] from Baker’s group used a graph based message passing neural network to autoregressively generate the protein sequences, and the sequence recovery is around 50% in the CATH dataset. Such a high sequence recovery rate makes it useful to design sequences for newly hallucinatory structures or *de novo* designed structures. However, the sequence diversity of most methods in the protein core region is not satisfactory and lacking side-chain atomic details hinder their applications. For example, in protein binder design using ProteinMPNN, the side chain packing of the initial guess, *i*.*e*. designed complex, is further modeled using Rosetta [29]. Inspired by AF2, McPartlon. et. al [30] recently developed AttnPacker to repack protein side chain with atomic coordinates, and predict the missing residue types for a given protein backbone. AttnPacker seems to give highly accurate side-chain conformation prediction, however, there is still a lack of experimental testing. In parallel with this work, Baker’s lab developed ligandMPNN [31], which could be used to predict the residue type and side-chain conformation under the condition of molecule context for molecule-binding protein design.

We consider that it is important for a protein sequence design algorithm to generate diversified sequences that can fold into the target structure than only presents native sequence-like ones, so that more can be learned about the sequence and structure relationship and new functions can also be easily introduced. Actually, Willson et al. [32] have shown that function is conserved for two proteins with about 40% sequence identity and the same fold. Todd et al. [33] have also demonstrated that when the pair-wise sequence identity is above 40%, over 90% enzymes have conserved function with the same four EC numbers. These analyses indicate that 40% of sequence identity might be enough to guarantee conserved function in natural proteins. Thus, pursuing high sequence recovery rates above 40% is not necessary and may be misleading.

Though deep learning based methods can implicitly learn high order interactions between protein residues using multiple neural network layers in the Encoder module, the Decoder module should be carefully designed to avoid label leakage of residue type in auto-regressive models, especially in the iterative models. For example, in the graph-based iterative model, in order to decoder the central residue type at node *i*, the node *i* aggregates the information of known neighboring residue node *j* and *k* in one GNN layer. Thus, the residue label may leak if we increase the GNN layer to decoder the residue type at node *j* as the information will flow from node *i* to *j*.

In order to leverage the power of GNN for both decodings of residue types and side-chain *χ*_1*−*4_ angles, and avoiding label leakage of residue types, we proposed GeoSeqBuilder, an algorithm to explicitly express the three-body interactions between residues in the Encoder module and thus each residue position could be trained in parallel without label leakage for fast inference during design. We employed a multi-scale graph convolutional network to learn the relationship between structures and sequences, which are used for learning different scale spatial local microenvironment of central residue, correspondingly. Considering the contextual dependencies and similarity to co-evolutional analysis, we utilized a conditional dependent scheme to capture the information from coupling interaction residues and generate novel sequences of the target structures. GeoSeqBuilder could recapitulate over 50% native residue types and simultaneously achieve decent side-chain conformation recovery. Direct coupling analysis showed that GeoSeqBuilder has learned the coupling relationships and is able to produce more diverse sequences relative to the references. We further utilized a variant network with shared input features as the sequence design module to specifically predict the side-chain conformation and obtained about 6% prediction accuracy improvement. Our learned likelihood confidence *logP* exhibited strong correlation with TM-score between protein decoys and native structures, which makes our algorithm versatile to discriminate the model quality during protein structure predictions and designs.

We have applied GeoSeqBuilder to design new sequences for two protein structures, including a commonly occurred natural thioredoxin fold, and a *de novo* hallucinated protein structure. The designed sequences could be expressed well in *E*.*coli* to give soluble and monomeric proteins with high thermal stability and correct secondary structures. The two solved high resolution crystal structures are in excellent agreement with the designed atomic models, and the other one solved crystal structure at low resolution demonstrates the good match of the protein backbones.

## Results

### Overview of model architecture

We devised a multi-scale graph convolutional network to learn the relationship between sequences and structures (Figure 1). Specifically, we use five-scale network to learn the spatial local microenvironment of the central residue where the 1-scale network is used to learn the two-body interactions, 2-scale network is used to learn the three-body interactions and i-scale network is used to learn i+1 -body interactions (Figure 1b), then iteratively determine the residue type and side-chain dihedral angles. To keep rotational and translational invariance of the protein, GeoSeqBuilder utilized local frames which are relatively geometric representations to extract the edges information (Figure 1c). GeoSeqBuilder takes full account of the coupling relationships between residues which is different from many previous studies that recover the central residue that fit to the environment without considering the coupling or dependence among residues (Figure 1d). To solve the problem of possible information loss by generating sequences with a fixed direction in auto-aggressive models, *i*.*e*. from the N-terminal to the C-terminal or *vice verse*, we also leverage the effects of later residues to better determine the favorable residue type in the current location (Figure 1e). We format the conception of the algorithm as:

**Figure 1.**
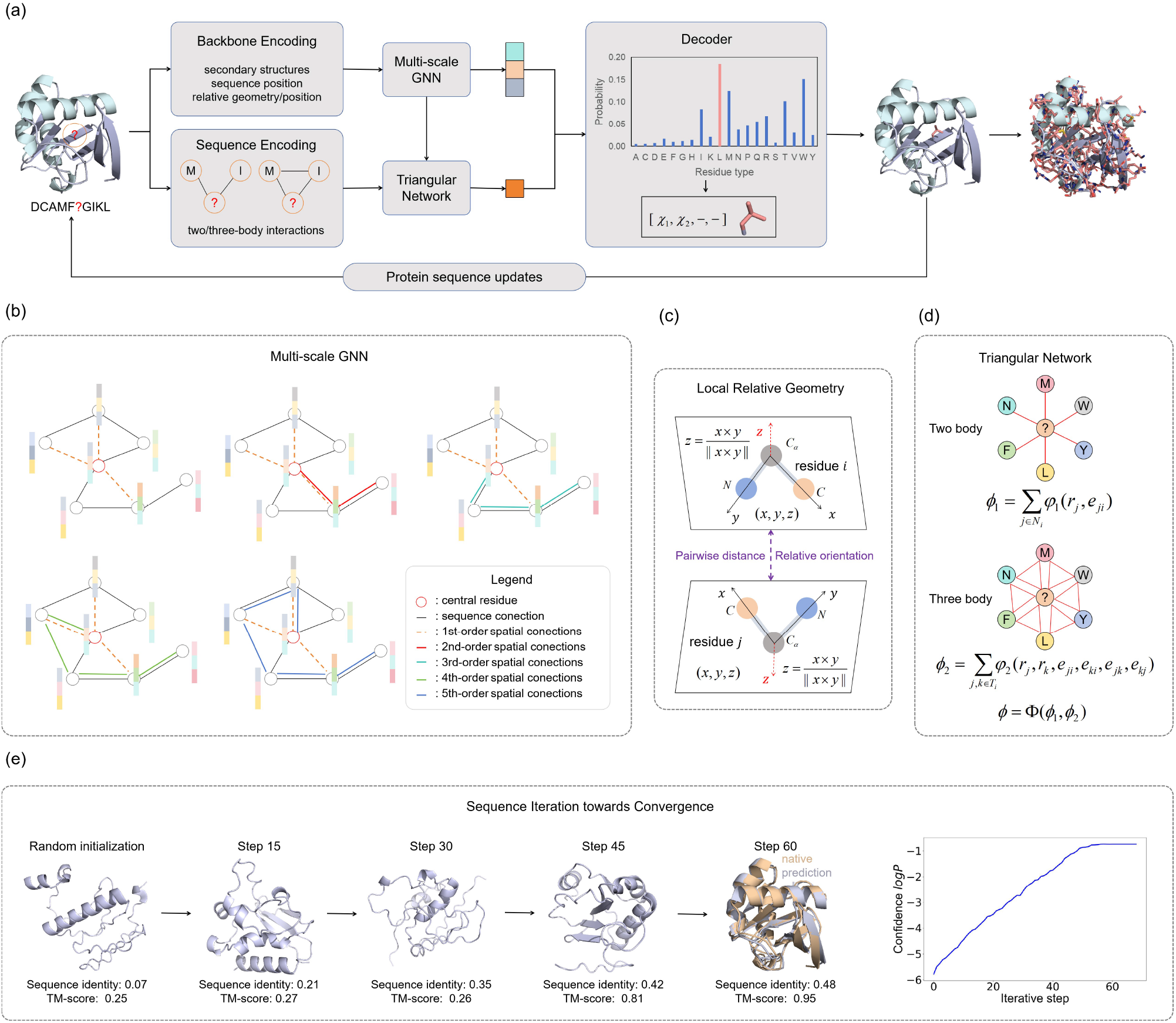
An overview of the protein sequence generative model used by GeoSeqBuilder. (a) A framework of Encoder-Decoder. In Encoder, all of the input features are from geometric properties of protein backbones. The multi-scale GNN modules are used for different-scale local information extraction. In Decoder, the self-masked scheme is used to learn the coupling relationships between other surrounding residues from local two-body and three-body interaction. The Decoder finally outputs the distribution of central residue type and the side-chain torsion angles which are sampled from the predicted distribution. The continuous side-chain torsion angles are transformed into 48 discrete bins (*i*.*e*. 7.5*°* a bin) from -180*°* to 180*°*. (b) Multi-scale network learns different visualizations from the surrounding paths. (c) The local frames are designed to extract the relatively geometric information from its spatially surrounding frames with translational and rotational invariance. (d) The triangular network is important to learn the two-body and three-body interactions. (e) During the design iteration process, The designed sequences are predicted using OmegaFold and the predicted structure of the end state is highly consistent with the native structure. After several circles of iteration, the predicted confidence log*P* reaches the convergent state.

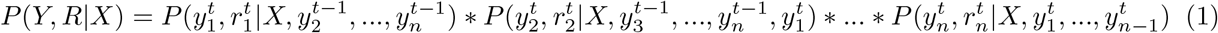

where *X* is the protein backbone, *Y* and *R* are the sequence and rotamer of comprised of *n* residues, correspondingly, and the 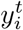 and 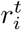 are the residue type and side-chain conformation of *i*^*th*^ at iteration *t*, respectively. In order to generate compatible sequences for the backbone, we trained the model by maximizing the likelihood function, and the *−logP* (*Y* |*X*) can be viewed as a scoring function to select the good generated sequences.

### Analyses of protein sequence recovery and properties

We trained GeoSeqBuilder using the CATH4.3 dataset [34] which contains 25k high-resolution single-chain protein structures from the Protein Data Bank and has been randomly split into training, validation and test set. See Methods for more details.

It is known that well-defined hydrophobic cores are essential for the stability of soluble proteins. As we have done in GeoPacker [35], we paid more attention to residues in the core region by employing normalized weight penalties from various locations to train the model,

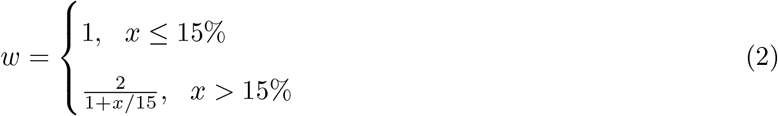

where *x* is the relative solvent accessibility surface area(rSASA). The residues with rSASA lower than 15% are defined as the buried while those with a higher rSASA are considered as exposed. The penalty for buried residues decreases as the rSASA value increases.

We evaluated several main influential encoding factors, and the corresponding results are presented in Figure 2a. Specifically, we add different side-chain pseudo atoms from CB to CG as most residues have CB and CG atoms,and we anticipate that encoding of incorporating those atoms could help algorithm to detect how much space remains or exceeds in that position. When we add pseudo CB atoms (Ala), the sequence recovery is between 46.83% and 48.63%. And when we further increase the pseudo CG1 and CG2 (Val), the sequence recovery increases to 49.24%. The possible interpretation is that inclusion of additive pseudo atoms CG1 and CG2 allows for a more comprehensive exploration of both sides of the cavity to enhance the accuracy of determining the suitable central residue types. We next add the triangular network (see Figure 1d and Methods) that further improve the quality of designed sequences as shown in the following tests. For residues with low abundance in native proteins including Cys, His, Met and Trp, the adjustment of slightly reweighting the label loss of those residues from 1 to 1.5 to make classification approximate balance for the 20 residue types further improve the predicted accuracy of central residue types to 51.58%. We also tested the influence of other settings, including different numbers of hidden dimensions, which did not cause much changes (Table S1). The learning curve is given in Figure S1. Overall, the model can recapitulate around 51% central residue types, which is similar to the recent experimentally validated works [27, 28]. Substitutions of the native residues with residues of similar physicochemical properties are observed (Figure 2b), for example, the Met can be replaced by Leu and Lys can be substituted by Arg. GeoSeqBuilder seems to give a higher propensity to use the negatively charged residues (Asp and Glu) than Asn and Gln compared to the natural proteins (Figure 2c). This observation can be further explored through the confusion matrix (Figure 2b). We further analyzed whether there are significant differences of the central residues recovery rate among different secondary structures verse the propensity of different residues varies across secondary structures. For example, Glu and Met are more inclined towards alpha helices, while Val is more prevalent in beta sheets. To this end, we calculated the predicted accuracy of individual residues according to their 3-state secondary structures (the accuracy of 8-state secondary structures is also available in Figure S2) and as expected, Tyr, Val, Ile, Thr, and Ser are predicted more accurately in beta sheets, while Glu and Met, and Asn are more accurate in helices and loops, respectively (Figure 2d). We further computed the predicted accuracy of individual residue types based on their rSASA. It is evident that hydrophobic residues in the core region are predicted more accurately than those exposed to solvent. This trend is also observed for the polar and even charged residues, which could be attributed to the additional constraints involved in forming satisfactory hydrogen bonds (Figure 2e). We also illustrated the correlation between rSASA and the confidence score of the central residue using entropy. The resulting curve can be approximated by a U-curve (Figure S3), indicating that while some exposed positions are more conservative, the majority of positions become more ambiguous as rSASA increases.

**Figure 2.**
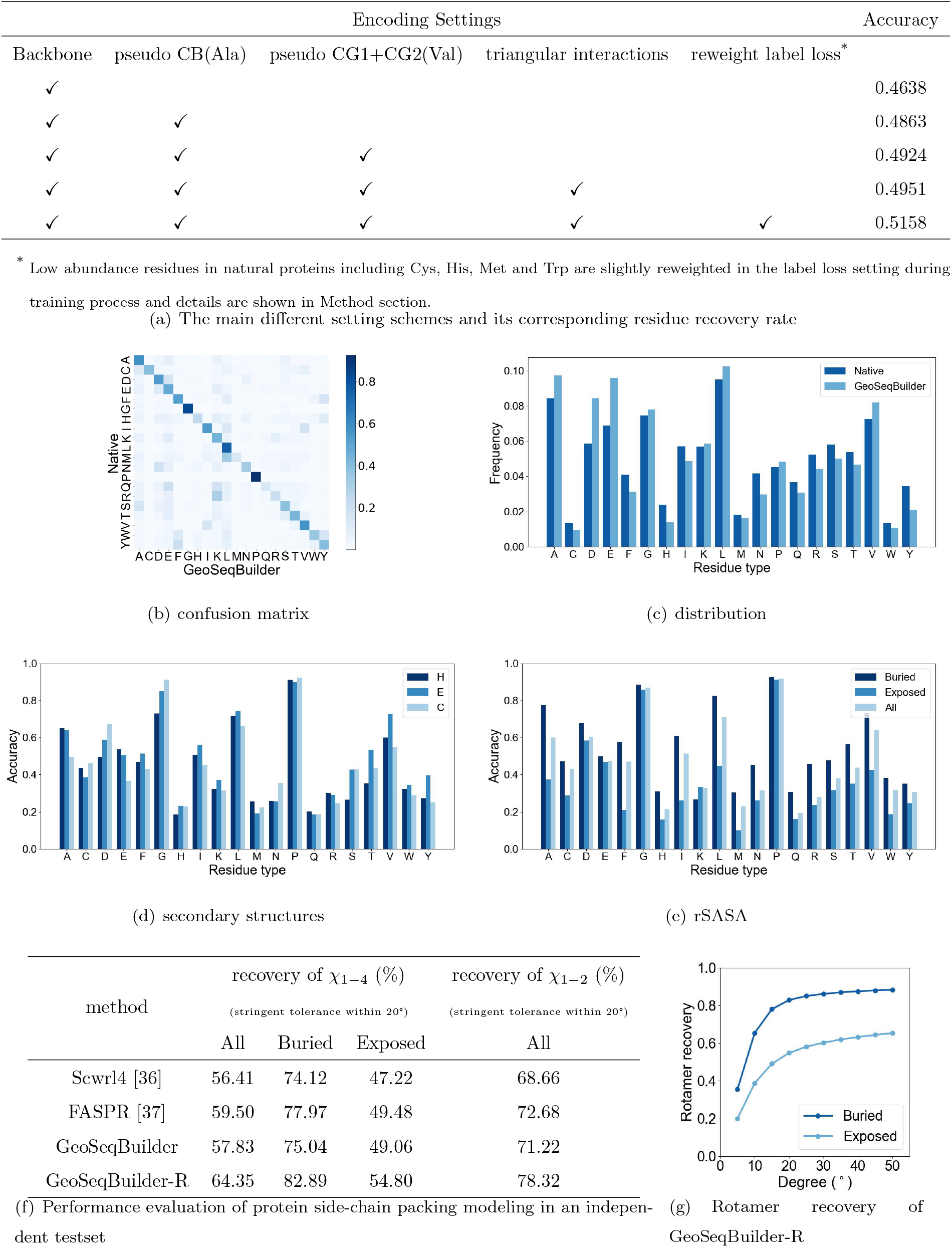
Performance of GeoSeqBuilder on an independent testset. a)Residue type recovery; b)The confusion matrix between the native and predictions; c) the individual residue frequency between the native and predictions; d) The predicted accuracy of central residue types for three secondary structures; e) The predicted accuracy of central residue types according to its rSASA; f) Protein side-chain conformation recovery; g) The results of GeoSeqBuilder-R on the independent testset for buried and exposed residues across at different error tolerance degrees.

Auto-regressive models generate central residue type one by one following a given sequential order, which may overlook the influence of later residues on the currently generated residues. The iterative algorithms may give convergent optimal central residue type. In response to this and to see how differences are between the generated sequences, we selected 41 proteins (see CAMEO41 for details) and generated 10 sequences for each backbone for evaluation. When starting from different initial states (*i*.*e*. the initial sequences are randomly given), we observed that the terminal convergent sequences for a protein target shared around 83% sequence identity with pair-wise sequence identities ranging from 60% to 90% depending on different targets (see Figure S4a). We found that the number of convergent steps is around a half of the sequence length of the given protein (see Figure S4b). It is interesting that the pair-wise sequence identity of the generated sequences for an engineered three-helix bundle protein ranges from 0.38 to 0.75 and all the predicted structures show high confident score with pLDDT exceeding 90 (see Figure S5 for the pair-wise identity matrix and some representative structure alignments). We also employed another scheme, starting from the native sequence, and the terminal sequences are more converged with around 0.9 sequence identity (Figure S6a).We further compared the results of starting from random sequences and from the native sequences. Apparently, random initialization gives more diversified sequences (Figure S6b).

In addition, we investigated the impact of sampling temperature on the diversity of generated sequences (detailed equation is given in Method). The sampling temperature equals 1 means that the sampling distribution is equivalent to the model output, and lower sampling temperatures result in sharper distributions. Our experiments revealed that there was no significant difference when the sampling temperature was below 1 (Figure S7), which may be attributed to the already sharp distribution of our predictions. To generate more diverse sequences, a plausible approach is to perturb the given backbone.

### Analyses of protein side-chain conformation prediction

For the first stage, our focus is primarily on the sequence design, which makes the side-chain conformation prediction not perfect. Specifically, we simultaneously predict the central residue type and side-chain torsion angles in an independent manner to accelerate inferring speed (see equation 3), which is different from the auto-regressive scheme. In the second stage, we utilize a triangular network with fewer parameters to refine the side-chain conformation prediction using equation 4. We incorporate the predicted central residue type and then predict its side-chain dihedral angles, which enhance the relationship between the residue type and its rotamer distribution in the backbone microenvironment. We next evaluate the performance using the independent test set and compared it with several conventional methods. We define the conformational recovery as only when all the predicted side-chain *χ*_*i*_ are within 20°to that of native structure. As depicted in Figure 2f, our algorithm demonstrates superior performance in recovering the *χ*_1*−*2_ at a stringent tolerance. However, conformation prediction of longer side chains needs to be improved. We then trained a variant network (See Method for details) with shared input features that predicts the side-chain conformation, namely GeoSeqBuilder-R, where ”R” signifies rotamers. As shown in Figure 2f, the side-chain conformation recovery of GeoSeqBuilder-R is more accurate than those predictions using GeoSeqBuilder, increasing by 6% in the large independent testset. The performance of GeoSeqBuilder-R is also significantly better than conventional side-chain packing approaches, including Scwrl4 [36] and FASPR [37]. To further inspect the degree of deviation to the native rotamers, we depicted the overall rotamer recovery of GeoSeqBuilder-R across different tolerance cutoff (Figure 2g). The rotamer recovery is rapidly improved when error tolerance increases from 5*°* to 20*°* not only for the buried residues, but also those exposed residues, and while the error is increasing after 20*°*, the rotamer recovery is gradually to a plateau, indicating at least one of side-chain torsion angles *χ*_*i*_ is remote from the natives.

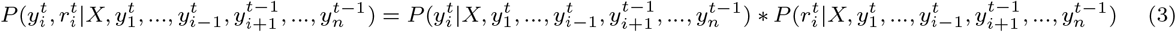

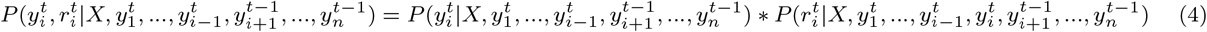

### GeoSeqBuilder learned the inter-residue coupling information

Inter-residues coupling analysis provides a better indicator to measure the backbone designability and quality of designed sequences compared to the sequence recovery rate. We employed two protein targets to evaluate whether our model learned the co-evolution information through direct coupling analysis(DCA) [38,39]. We utilized our pre-trained model to generate 200 sequences starting from random initialization for the two protein backbones, including an alpha-helix bundle (PDB entry: 7DGU) and thioredoxin (PDB entry: 1FB0). We further employed Rosetta to perturb the native backbone [40], and generated 50 decoys for each protein with root mean square deviation (RMSD) values ranging from 1.2Å to 1.7Å compared to the native structures. Our analysis revealed that slight perturbations to backbone lead to the generation of more diverse sequences (Figure 3c) as the perturbations make the backbone more tolerant to changes of residues which carries more co-evolution information (Figure 3a). However, perturbing the helix bundle protein 7DGU did not significantly improve the sequence diversity (Figure 3d), which might be caused by the inherently large sequence space available for helices. The relationship between DCA values and distance could be seen in Figure S8a-S8b, and indeed, the region of large values of DCA are located in the area of inter-residue contacts. The sequence logos of both the native and the perturbed structures are shown in Figure S8c-S8f.

**Figure 3.**
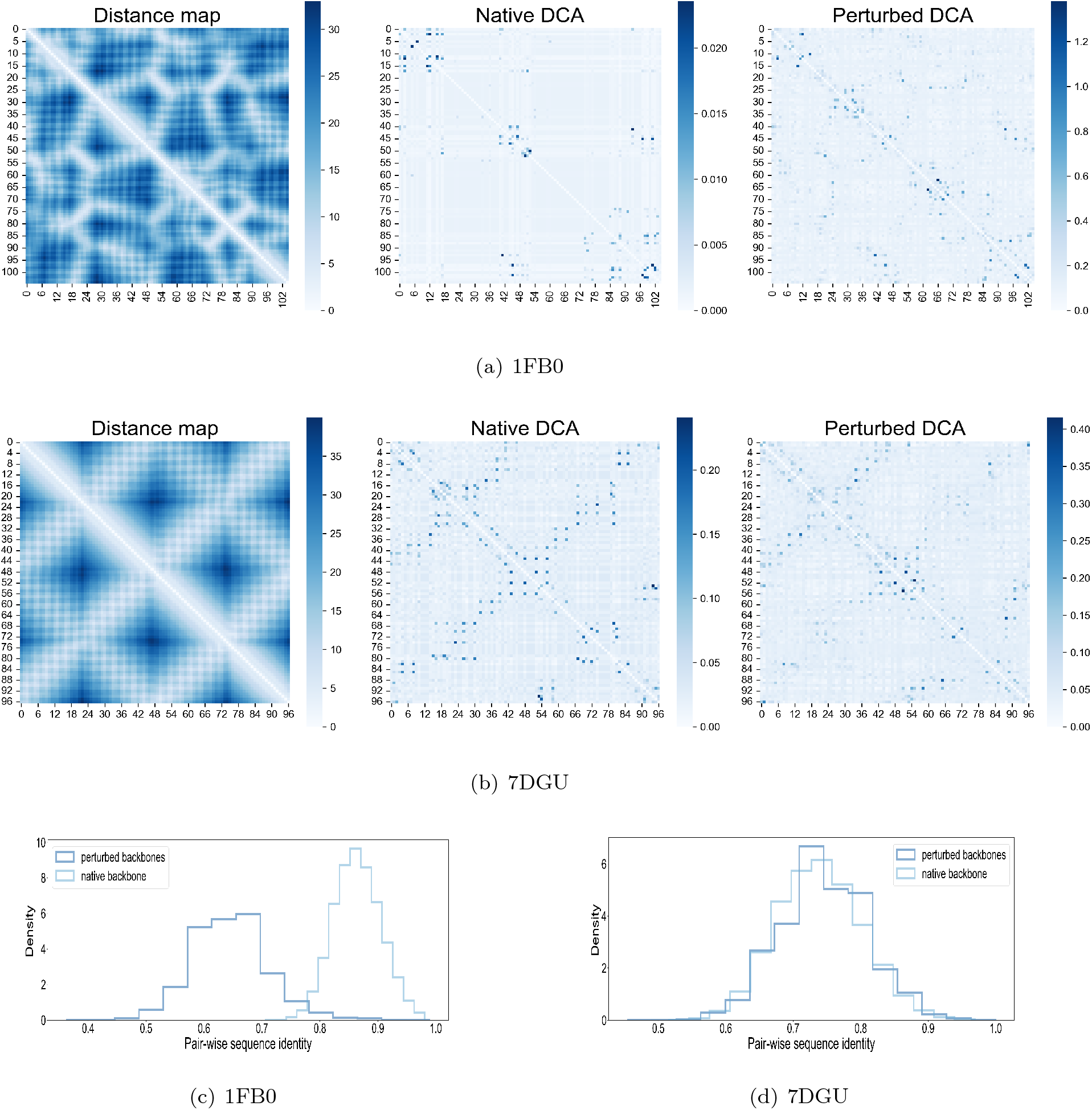
Direct coupling analysis for the native backbones and perturbed backbones. (a) The properties for protein 1FB0. From left to right column: distance map, the DCA plot from 200 designs for the backbone 1FB0 and the overall DCA plot from 200 designs for all 50 perturbed backbones.(b)The properties for protein 7DGU. (c)The distribution of generated pair-wise sequence identity for the native backbone 1FB0 and the perturbed structures. (d) The distribution of generated pair-wise sequence identity for the native backbone 7DGU and the perturbed structures.

### Scoring function evaluation on three main topological backbones

In order to assess whether the learned likelihood *logP* (*Y* |*X*) could be interpreted as a pseudo potential function, we conducted tests on three proteins, spanning from three main topologies-all alpha, alpha-beta, and all–beta, which are randomly selected from the testset. For each backbone, we started from a random sequence, and then iteratively updated the non-optimal central residues one by one until convergence. The resulting generated sequence identity relative to its native sequence is from 30% to 44%. In the process, the generated sequences are repacked using GeoSeqBuilder-R and energy minimization is further performed to eliminate the clashes of atoms using AMBER. Intuitively, if the likelihood *logP* (*Y* |*X*) performed well, we can expect it to exhibit a strong negative correlation with the Rosetta energy score function ref2015 [41]. Therefore, we utilized ref2015 to score each intermediate structure obtained through the process above. Notedly, all the three trajectories displayed the expectant tendency. Furthermore, the convergent sequences in all cases exhibited lower or similar Rosetta energy score compared to their native sequences (−262.352 vs -178.210 for backbone 1elk, -206.212 vs -200.250 for backbone 4ggt, and -147.396 vs -76.380 for backbone 4efo). To further validate the convergence sequences, we employed single sequence based AF2 for structure predictions. Interestingly, the AF2 successfully predicted the designed sequences, but failed to give correct folds when only the native sequences were used. Essentially, AF2 is multiple sequence alignment (MSA)-dependent algorithm for protein structure prediction, and its performance might be limited for single-sequence prediction. As pre-trained protein language models develop rapidly, the representative models, such as trRosettaX-Single [42], ESMFold [43] and OmegaFold [44] are promising and could be valuable for single-sequence based structure prediction. Therefore, unless otherwise specified, we later use OmegaFold to predict the structures of our designed sequences.

### The likelihood confidence *logP* (*Y* |*X*) is a good indicator to recognize the high quality protein decoys

Protein model quality assessment is a significant part of protein structure prediction, which aims to score the quality of protein decoys derived from various structure modeling tools, while there is no information about the ground truth [45]. Our algorithm of backbone-fixed sequence design learned the compatibility between sequence and structure based on the natives, thus, it may provide an alternative way to evaluate the quality of those decoys using our algorithm where the two hypotheses are that the designed sequences for the high quality decoys will lead to larger likelihood *logP* while the designs based on the low quality decoys will produce lower likelihood *logP*, and the similar sequences share the similar structures [46]. To this end, we used CD-Hit [47] to remove the redundant data in DeepAccNet [48] that share sequence identity higher than 40% compared to our training set. The dataset collected by DeepAccNet is non-redundant with pair-wise sequence identity lower than 40, and around 150 decoys have been produced for each native protein, which are derived from three methods, including native structure perturbation, comparative modeling, and deep learning guided folding. We randomly choose 50 targets as well as their decoys and designed 10 sequences for the protein backbones. Due to the subtle difference of likelihood *logP* within a backbone, we utilized the average likelihood *logP* as a score to evaluate the quality of backbone. We observed that there is a strong correlation between the likelihood *logP* and TM-score that the average Pearson correlation coefficient reached 0.65. The overall correlation distribution is shown in Figure 4d and four representatives are shown in Figure 4e. All of the correlations of those 50 proteins can be found in Figure S9.

**Figure 4.**
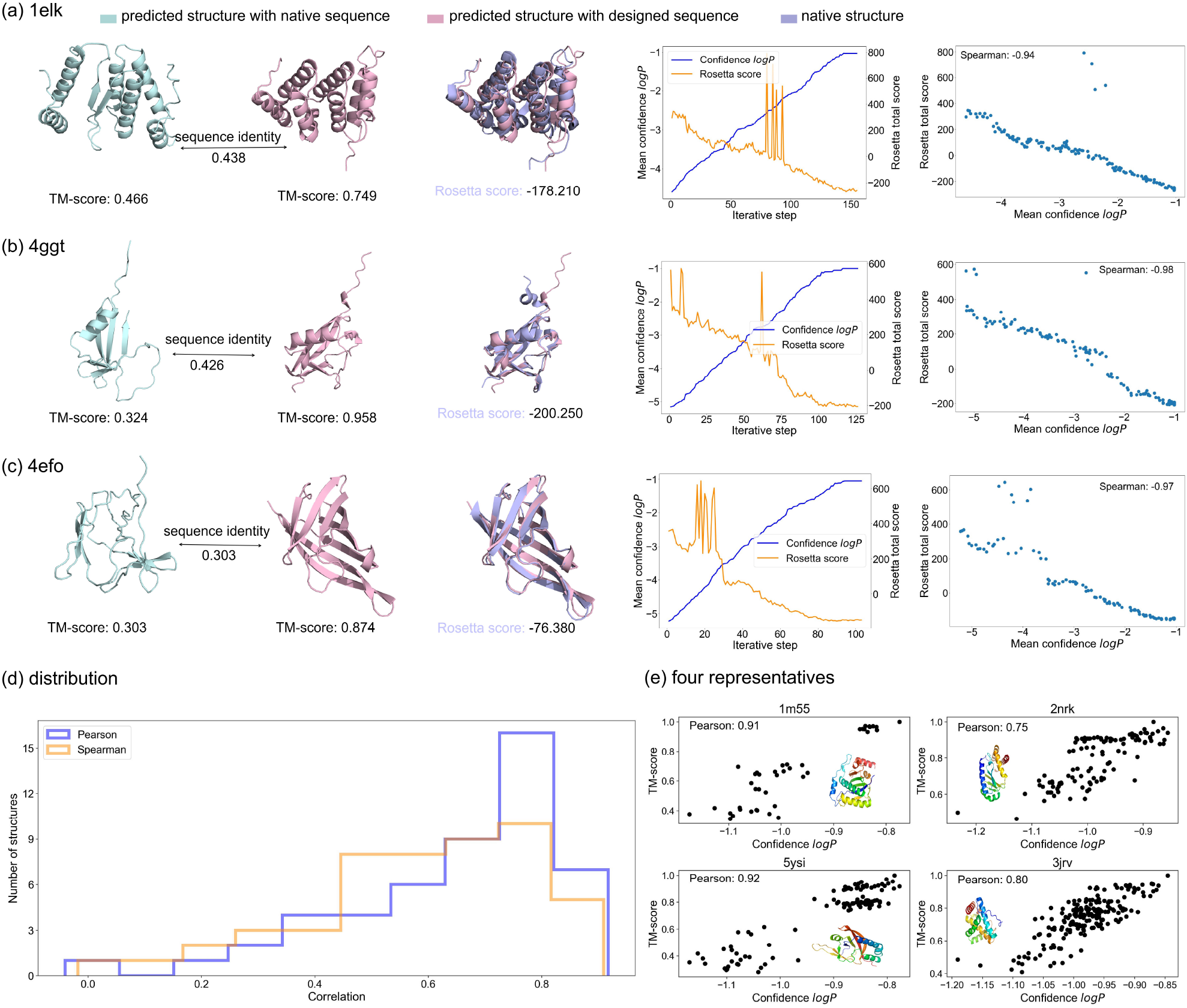
GeoSeqbuilder can design highly confident sequences on three main topological structures using AF2 metrics and pick up the near-native structures from decoys pool. (a) From left to right: the columns are the predicted structure of native sequences, the predicted structures of designed sequence, structural alignment between native and prediction, scoring metrics for a sequence sampling trajectory using GeoSeqBuilder and Rosetta, and correlation between Rosetta score and confidence logP of GeoSeqBuilder, correspondingly. All of those predicted structures are using AF2 in single sequence mode. (b) and (c) are similar presentations with (a). (d) The logP values of GeoSeqBuilder exhibit strong correlation to TM-scores between decoys to that native structure, and that the correlation distribution of 50 test structures demonstrates that the most are around 0.8. (e) Four representatives are listed to see how is the correlation.

### Model evaluation on natural and *de novo* designed structures for sequence redesigns

In order to further evaluate our model’s performance on the native proteins and compare it with other related methods, we utilized the recently released (between 2022-12-09 and 2023-06-03) monomer proteins from CAMEO [49] which are at high resolution (*≤*3Å) and without chain breaks, including 1 hard targets and 40 medium difficulty target as those native proteins are predicted with relatively low pLDDT (see Table S2 for details for those PDB entries). This choice of data allows for a fair comparison, as none of the methods used for comparison have encountered these specific proteins. As the Figure 5a shows that incorporating the term of explicit three-body interactions definitely empower the performance in all metrics, including pLDDT, TM-score, exposed hydrophobics. In general, the pLDDT shows the confidence of predicted model, and the TM-score shows how structurally similar is between the predicted model and the desired model. Besides that, the exposed hydrophobics is expected to be as small because it is prone to aggregation as many hydrophobic residues are exposed to solution.

**Figure 5.**
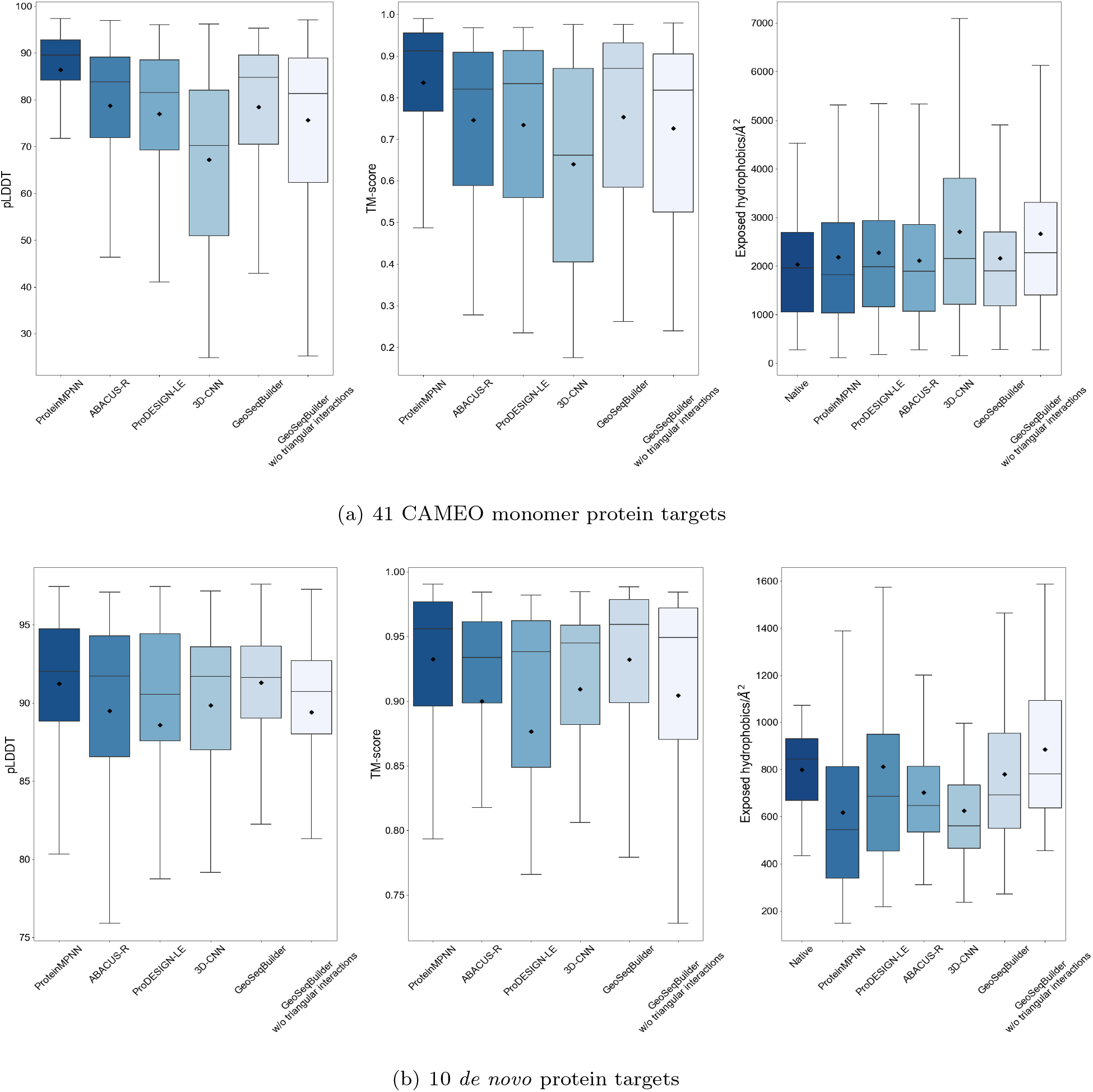
Metrics evaluation of the designed sequences using OmegaFold on two protein datasets. (a) The distribution of pLDDT, TM-score and exposed hydrophobics of the generated sequences for CAMEO41. (b) The distribution of pLDDT, TM-score and exposed hydrophobics of the generated sequences for the 10 *de novo* designed structures.

We generated 10 sequences for each protein target and used OmegaFold to predict the corresponding structures. Although incorporating the term of three-body interactions did not result in significant gains in sequence recovery, the increases in average pLDDT (3.7%) and TM-score (3.8%) are remarkable. Additionally, improvements were observed in the metrics of exposed hydrophobics. We further compared GeoSeqBuilder with representative deep learning sequence design approaches, including graph based ProteinMPNN [28], 3D-CNN based method [22], and transformer-based ABACUS-R [27] and ProDESIGN-LE [50]. The average pLDDT and TM-score (78.42 and 0.75) of GeoSeqBuilder were similar to those of ABACUS-R (78.76 and 0.75) and ProDesign-LE (76.99 and 0.73), better than 3D-CNN (67.19 and 0.64) and worse than ProteinMPNN (86.42 and 0.84). Besides that, the pLDDT and TM-score of GeoSeqBuilder were more concentrated in the upper median values compared with the ProDESIGN-LE and ABACUS-R. We also compared the exposed hydrophobics and observed that all methods, except 3D-CNN, exhibited similar values to the native proteins. We also compared the sequence recovery across all methods, which is depicted in Figure S10.

The aforementioned analysis demonstrates that GeoSeqBuilder is capable of generating new sequences that fold correctly to the desired structure for natural structures. The emergence of hallucination and diffusion models [51–54], generate possible *de novo* protein structures. Here we further investigate whether GeoSeqBuilder can generate plausible sequences for the previously unseen structures. To this end, we further assessed and compared the performance with related methods on recently released 10 *de novo* proteins (Table S3), including 4 mainly alpha, 3 mainly beta and 3 alpha beta structures. Similar to our previous findings, incorporating the term of three body interaction contributes to the quality improvement. The average pLDDT and TM-score (91.31 and 0.93) of GeoSeqBuilder are comparable to ProteinMPNN (91.24 and 0.93), and better than other methods (Figure 5b). It is worth noting that the predicted quality of all the designed sequences for the *de novo* protein structures shows significant improvements compared to the results above for natural protein structures, probably due to the more regular backbones used in *de novo* generated structures.

### Experimental validation on two protein structures

We further tested GeoSeqBuilder on two protein structures, including one natural protein structure and one hallucinated structure for sequence design. The first target is a thioredoxin fold [55] (PDB entry: 1FB0) which is prevalent in many species. The second target is 0705, a previously published hallucinated protein structure using trRosetta [51], for which no mono-disperse proteins and crystal structure were obtained before.

We first generate 200 sequences for each protein backbone structure and then cluster them using CD-Hit and analyze the sub-clusters to select the most diverse representative sequences. These selected sequences are without any artificial modifications and further undergo visual inspection using pymol at the atomic level as our model generates full atomic protein structures. We selected 9 and 6 sequences for 1FB0 and 0705, respectively for experimental testing. The sequence identities of the selected sequences for 1FB0 to the wild-type sequences are around 50%, while that of 0705 is as low as 23% (Figure S11). We further searched the non-redundant sequence database from the high quality clust30 [56] using the query sequences from 0705 designs, and found no homologous sequence, indicating that those sequences are indeed *de novo* designed.

We expressed the designed sequences in *E*.*coli*. All of the 9 sequences for 1FB0 and the 6 sequences for 0705 can be highly expressed and give soluble proteins. We purified all the 15 soluble proteins (Figure S12) and verified their monomeric state using gel filtration chromatography (Figure 6 a-c and Figure S14). We next examined the thermal stability of the proteins using nano Differential Scanning Fluorimetry (NanoDSF) and analyzed their secondary structures using circular dichroism (CD). All the designs for 1FB0 showed high thermal stability with melting temperature (Tm) higher than 110*°*C (Figure S15), significantly surpassing that of the wild type protein (68*°*C). Similar phenomena are observed for 0705 (Figure S15). CD spectra demonstrate that these proteins have expected secondary structures consistent to the target structures (Figure S16).

**Figure 6.**
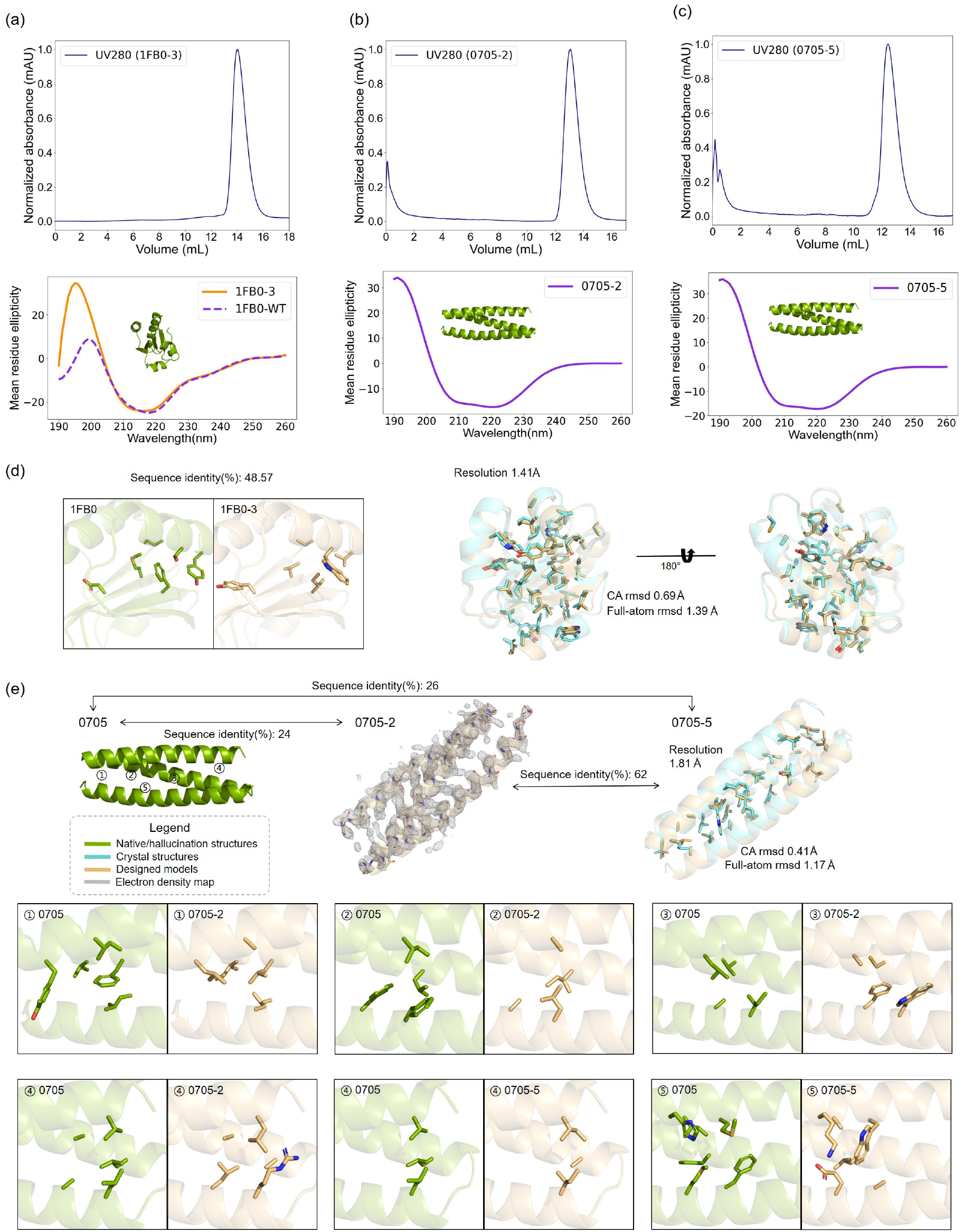
GeoSeqBuilder successfully designed sequences compatible with the target structures. (a) The experimental results of 1FB0-3. From top to bottom: the rows are results of molecular exclusion chromatography and CD wavelength scan, correspondingly. (b) and (c) are in similar presentation with (a). (d) The obtained high quality crystal structure 1FB0-3 shows different core packing with the native, and the atomic differences between the designed and crystal structure. (e) Multi different core packing from the crystal structure 0705-2 and 0705-5. 0705-2 only shows the electron density map and matched protein backbone without atomic details for the low resolution at 2.99Å. The different details of circled number are enlarged for comparisons of residue types.

We further performed crystallization conditions screening and successfully obtained 4 protein crystal hits, including 2 designs for 1FB0 and 2 designs for 0705, and optimized 3 of them for data collection (Table S4). We solved one structure for 1FB0 design and two for 0705 designs by molecular replacement using the designed sequence and structure models as input. As for the design 1FB0-3, we obtained the crystal structure at high resolution (1.41Å) and the overall protein structure superimposes with the design model very well with an RMSD of 0.69Å for all the *C*_*α*_ atoms (Figure 6d). The RMSD for buried side chains is 0.79Å and that for exposed side chain is 1.62Å. The sequence alignment reveals that 1FB0-3 shares overall 48.57% sequence identity to the natural sequence. Specifically, the buried residues and exposed residues are with identity of 72.97% and 35.29%, respectively. More careful inspection finds that the core triangular residues of 29M,58I,85F are changed to 29V,58V,85I (Figure 6d). For the designs based on the hallucination structure 0705, we solved structures for 0705-2 and 0705-5 at 2.99Å and 1.81Å resolution, respectively. Although we could not compare the details of the structures due to the limited resolution and low quality of diffraction data, the overall backbone structure of 0705-2 is consistent with the designed protein backbone, indicating that the designed sequence is folded correctly. The sequence of 0705-2 shared an overall sequence identity of 24% to 0705, and that the side-chain residue types in the core region are almost completely changed with 33.33% sequence identity to that 0705. Some of those residue rearrangements in core regions are exhibited in Figure 6e. The crystal structure 0705-5 at 1.81Å clearly shows side chain interactions and the core region is also repacked differently from the hallucination structure 0705 and crystal structure 0705-2 (Figure 6e). 0705-5 shares an overall sequence identity of 26% and core sequence identity of 50% and 73% to 0705 and 0705-2, respectively. The RMSD of 0705-5 designed model and crystal structure is 0.41Å for all *C*_*α*_, 1.17Å for full atoms, 0.39Å for the buried side chains and 1.28Å for the exposed side chains, respectively. The details of crystal structures are shown in Figure 6. The two high quality crystal structures have been deposited in the Protein Data Bank (1FB0-3 for PDB entry 8XYW and 0705-5 for PDB entry 8XYV). The experimental investigation demonstrates that GeoSeqBuilder can successfully design well-folded sequences with abundant diversity of core packing arrangements.

## Discussion

We have employed a multi-scale graph convolutional network, GeoSeqBuilder, to iteratively decode the central residue types and side-chain torsion angles for structure guided protein sequence design. Compared to the state-of-the-art methods, ours model exhibits several novelties: firstly, we employed the pseudo CG1 and CG2 to explore the cavity while ABACUS-R and ProteinMPNN utilized the backbone and pseudo Ala; secondly, our model is an all-atom model that gives the side-chain atomic coordinates which is useful to study the atomic interactions especially in the scenario that the predicted models are unavailable or wrong; thirdly, our algorithm can recognize high quality protein structure decoys, which is an alternative approach to evaluate the quality of structure decoys. In addition, we also find more diversified sequences with high confidence scores can be generated by slight backbone perturbation.

We experimental tested the design ability of GeoSeqBuilder using two protein structures. All of the 15 sequences designed without any post modifications by human expertise are soluble and highly thermal stable. The two high resolution crystal structures reveal structures that are consistent with our designed atomic models, especially in the core region. The generated atomic models give more information for inspection and do not need protein structure prediction using other tools as most of the other methods do. Atomic model is also useful in protein-protein interaction design and molecule-binding protein design, as most of those designed interactions are contributed by protein side chains. We further trained a variant network with shared input features of GeoSeqBuilder to increase the side chain conformation prediction accuracy, the resulting GeoSeqBuilder-R has been integrated into one package as user-friendly tool, namely GeoSeqBuilder.

More careful inspections reveal that GeoSeqBuilder can generate well-defined diverse core residue rearrangements, which is useful to find new sequences and functions during protein design. For example, the designed sequences for 0705 are significantly different from the hallucination sequence using trRosetta and remote from the natural sequence space. Most triangular residues in the core region are repacked which might be caused by introducing the triangular network to learn the three-body interactions (Figure 6d-e).

In backbone-fixed *de novo* protein design, sequence diversity plays a crucial role. The ability to explore a wide range of sequence space is essential for discovering different core packing arrangements, which can lead to the identification of novel and stable protein folds. However, it is important to strike a balance between sequence diversity and the success rate of the design process. Excessively high sequence diversity can pose challenges in the design process. It may lead to a larger number of non-functional or unstable sequences that fail to fold correctly or adopt the desired structure. This can result in a lower success rate and increase the difficulty of identifying viable designs. Balancing sequence diversity is, therefore, a significant challenge in *de novo* protein design. Besides that, the supervised learning also limits the generated sequence diversity. In other words, a protein backbone should be corresponding to many similar sequences in natural proteins, however, this natural phenomenon has not been fully considered. In this study, we found that slight perturbations of the protein backbone can increase the sequence diversity. The high correlation between our confidence log*P* of model and TM-score of decoys of native structure provide further support and criteria for the extent of tolerant perturbations. To further enhance sequence diversity, if more than one structures (*e*.*g*. in case the one protein target binds to different ligands or the NMR structures) are available for the target protein, multiple structures can be used for protein design. The other approach is to employ a multi-stage generation strategy. In the initial stage, GeoSeqBuilder is utilized to generate atomic models based on the provided backbone, followed by energy minimization, or even molecular dynamics simulations of these models. Subsequently, in the subsequent stages, the energy-minimized models can be used as backbones and fed into GeoSeqBuilder for further rounds of model generation. By repeating this process for several rounds, we anticipate that GeoSeqBuilder will be capable of generating sequences with remarkable diversity.

One of the limitations that cannot be overlooked in the current version of GeoSeqBuilder is the absence of encoded information regarding small molecules, nucleotides, and metal ions. This omission has a notable impact on the success rate of protein designs, particularly in cases involving molecule binding. The design of proteins that interact with specific molecules heavily relies on the accurate representation of these molecules and their interactions. Without encoding relevant information, GeoSeqBuilder is unable to capture the intricacies of these interactions, leading to suboptimal designs and reduced success rates. Introducing the encoding of small molecules, nucleotides, and metal ions into GeoSeqBuilder would be useful for molecule-binding protein design, and as a priority, we anticipate that GeoSeqBuilder could accurately exhibit the atomic interactions between protein side chains and molecules. This would be done in the future version.

## Methods

### Algorithm

Our model, incorporates two pipelines, one for protein backbone graph encoding, and one for iteration recycling of sequence encoding using triangular network, which is shown in Figure 1.

In the GNN module, the protein chain is regarded as a map, and each residue is at a node, while the edge is used to connect these nodes based on a given distance threshold between two *C*_*α*_s. In the node level, the features include backbone torsion angles (ϕ,ψ) as well as its 4-order Fourier series expansion, and one-hot encoding of 8-state secondary structure elements derived from DSSP. Besides, the degree of nodes reflects the properties of a map, and thus the degree of a node and its inverse are both added. After that, the node features are totally with 26 elements.

If the distance between two distinct *C*_*α*_s is within 12 Å, a link is built. A local frame of central residue is comprised of three backbone atoms (CA,C,N) where the origin is located at *C*_*α*_ coordinate, and the *C*_*α*_ *− C* and and *C*_*α*_ *− N* is along with X-axis and Y-axis, respectively, and Z-axis is orthogonal with the plane of C-CA-N. The relative orientation of counterpart atoms of surrounding residues within 12 Å can be both represented in a tuple with 3 elements. In this manner, the extracted features keep rotation and transformation invariance, which is very significant within the graph encoding. The distance between two atoms are further expanded to its power series form 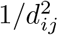 to 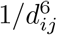 plus its original value *d*_*ij*_. Thus the edge features are comprised of 343 elements by pairing total atoms within corresponding residue pairs. Besides that, the information of relative location of residues in sequence is encoded into the edge using the difference between *i* and *j* with cosine and sine value. Thus, a vector with 345 element is mapped to the terminal edge representation.

For the embedding of node features and edge attributions, we firstly use a linear layer with elu activation, then those hidden layers of nodes and edges are multiplied to preserve the dependency between those spatially connected node and edges. The underling equations (5) and (6) provide a concise explanation of information updates of nodes and edges.

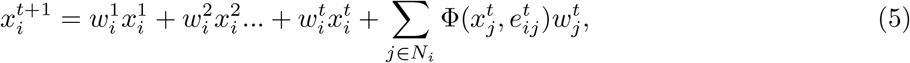

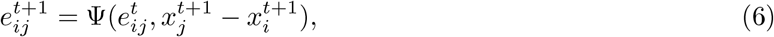

where 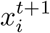 denotes the iteration t+1 of node *i, N*_*i*_ is the set of neighbors of node *i, w*_*i*_ is a learnable parameters which reflects the importance of residues, Φ, Ψ are both learnable functions and *e*_*ij*_ is the edge. Here Φ, Ψ are linear with activation function *elu*. An tuition is that the node information are from the previous node message and are gathered along with the current neighbors edge attributions.

This ensures that the node embeddings capture relevant information from both the node itself and its neighboring nodes. As for the edge updates, we incorporate the difference between node *x*_*i*_ and node *x*_*j*_ to facilitate the information flow. This allows the model to learn and incorporate the relevant information between connected nodes. Finally, we employ a 5-scale GNN architecture, where each scale GNN follows the same node and features update process described above. This multi-scale approach enables the model to capture and leverage information at different levels of granularity within the graph structure.

In our approach, the sequence residue type is encoded as a 20-element vector. After processing the protein graph map, the residue coupling relationship is further learned with a diagonal masked matrix in the decoder module, where the diagonal elements are set to 0 while the rest are 1. This design allows central residue to have a broad view of all the other surrounding residue types, except for itself. This information plays a crucial role in capturing two-body interactions between residues. Besides that, triangular interactions where these distances are both within 12Å are incorporated. This is motivated by the observation that two residues are often influenced by another residue within a triangular arrangement, thus two-body interactions alone are not sufficient to capture the relationships. To address this, we employ a triangular network to learn the the two-body interactions and triangular interactions along with the embed graph information from backbone Encoder module. Those detailed triangular network has been illustrated in Figure 1d. The output of high-order information of residue-residue interactions from triangular network and multi-scale network are further integrated and fed into the MLP (*i*.*e*. Decoder module) to determine the terminal central residue type and simultaneously predict the side-chain dihedral angles in an independent manner. More specifically, The output logits of Decoder module represent the distribution of central residues with 20 elements and 4 side-chain dihedral angles *χ*_1*−*4_ with (4,49) elements, correspondingly. The side-chain torsion angles, from *χ*_1_ to *χ*_4_ are split into 48 bins (*i*.*e*. 7.5° a bin) from -180° to 180° plus an additional bin for the BLANK state. For example, the *χ*_4_ for Met is BLANK. Usually, the central residue type with predicted maximal probability is expected to be consistent with the native one.

These details and explanations have been elaborately described in Figure S17 which serve as the supplementary material for Figure 1 in this paper.

The Loss function used in training includes two parts:

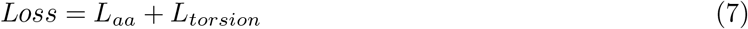

where the first term is the residue type loss, the second is side-chain torsion angle loss. Both terms are measured by the cross-entropy function and normalized by the transformation of rSASA mentioned above. The weight penalties of these residues with lower frequency in native proteins, including Cys, His, Met, and Trp, are slightly changed from 1 to 1.5, 1.5, 2 and 2, correspondingly. All of these penalty values are empirical settings without many efforts for tuning.

The sampling temperature *t* controls the sequence diversity in a position using the following equation:

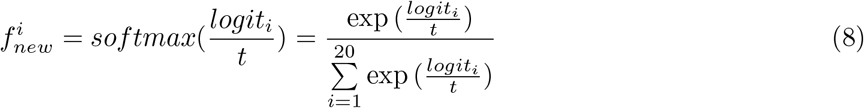

where the *logit*_*i*_ is the model output of *i*^*th*^ residue type in a position. We highly recommend users to utilize the default sampling temperature of 0.1 during their experiments. This temperature setting strikes a balance between efficiency and generating sequences with moderate diversity for comprehensive test verification. On the other hand, we do not recommend using sampling temperatures larger than 1. Higher temperatures smooth out the sharpness of the predicted distribution, resulting in sequences that may lack specificity. Moreover, higher temperatures can increase the number of convergent steps in the sampling process, potentially leading to the generation of inferior sequences.

The variant network GeoSeqbuilder-R was developed to specifically predict the side-chain torsion angles. As a result, there are some key differences in the network architecture compared to GeoSeqBuilder. In GeoSeqbuilder-R, the sequence information is merged with the node features instead of using a separate triangular network for sequence encoding. This modification allows for a more direct integration of the sequence information into the node features, facilitating the prediction of side-chain torsion angles. Furthermore, GeoSeqbuilder-R focuses solely on predicting the distribution of the four side-chain dihedral angles *χ*_1*−*4_. It does not involve predicting the distribution of central residues or other aspects of protein structure. By specializing in side-chain torsion angle prediction, GeoSeqbuilder-R is optimized for tasks that require accurate modeling of side-chain conformations and can be particularly useful in applications such as protein structure prediction and protein design. Integrating GeoSeqBuilder-R into GeoSeqBuilder for atomic protein design, users can both combine the strengths of both models, and improve the computational efficiency and inference speed for the shared input features when compared with two separate algorithms.

### Data

We use a high resolution X-ray crystal structural dataset (*≤*2.5Å) from PDB based on the level of Orthogous Seqeuce Family(S60) at recently released CATH4.3, as well as protein chain length in the range of 50 to 600. The NMR structures and membrane proteins in S60 are eliminated due to the flexibility and specificity, respectively. In nature selection, some accepted substitutions are often appeared in a protein families and cause the subtile perturbation of protein structure. Thus, the non-redundant data S60 could provide some of those useful information, which are captured by our model. Finally, a set with total 25067 proteins is randomly split into a training set containing 23500 proteins, a validation set including 567 proteins and a remaining independent test set comprised of 1000 proteins.

### Implementation and runtime

The algorithm has been implemented by Python. The network architectures are built under the framework of PyTorch and PyG libraries. The maximal number of network layer(*i*.*e. S*_*N*_) and hidden unit, and the radius of contextual environment have been explored, and finally, these values have been set to 5, 50 and 12, correspondingly. Each layer module is comprised of a linear transformation, batch norm and an elu activation function. In the last multi-perception layer, a dropout rate 0.5 is used for avoiding overfitting. The optimizer is ADAM, and the learning rate is set to 0.001. After about 15 epoches, the train validation reaches the optimal state. It spends around 2 hours for each epoch during training on a single A30. Our inference is much faster, and it spends around 2s to obtain a convergent sequence for a protein length of 100. Beginning from a random sequence, the convergent iteration steps equal to around a half of protein sequence length.

### Experiments

The DNA sequences of the designed proteins were codon-optimized and synthesized by Beijing SYKM Gene Biotechnology Company, and cloned into expressing vector pET45b and pET28a for 1FB0 and 0705, respectively. The plasmids were transformed into *E*.*coli* BL21 (DE3), and the bacteria were cultivated in the lysogeny broth culture medium with 1mM Kanamycin. 0.5mM IPTG was added when the *OD*_600_ value is between 0.6 and 0.8 to induce protein expression for 15 hours at 18*°*C. These bacteria were further sonicated in buffer including 20mM PBS and 500mM NaCl at pH 7.6. The soluble supernatant was purified by Ni2+ affinity chromatography and eluted at solution containing 20mM PBS, 500mM NaCl, and 500mM Imidazole. These initially purified proteins were further isolated using Superdex75 10/300 GL column in buffer containing 20 mM Tris and 300 mM NaCl at pH 7.3. The concentration of protein with Trps was directly measured using NanoDrop while those without Trps are determined using standard BCA protocol and performing 3 times parallel gradient assays. The melting temperature for characterization of protein thermal stability were done using NanoDSF (NanoTemper) with protein concentration at 0.2mg/ml. Proteins for CD characterization were transfer to the solution containing 20 mM KH2PO4/K2HPO4 at pH 7.4 and concentrated at 0.1-0.2 mg/ml, and data were collected on a MOS-450 AF/AF-CD spectropolarimeter (Bio-Logic). The crystallization conditions were listed in Table S4.

## Abbreviations

RMSD: Root Mean Square Deviation
pLDDT: predicted Local Distance Difference Test
NanoDSF: nano Differential Scanning Fluorimetry
CD: circular dichroism
Tm: melting temperature.

## Ethics approval and consent to participate

No applicable.

## Consent for publication

No applicable.

## Availability of data and materials

The source code, along with the pre-trained model will be available in https://github.com/PKUliujl/GeoSeqBuilder. The solved crystal structures have been deposited in the Protein Data Bank with the accession codes 8XYW (1FB0-3) and 8XYV (0705-5).

## Authors’ contributions

LL and CZ conceived the project and computational framework. JL performed the computational and experimental study. ZG collected the crystallographic data and solved the crystal structures. LL, CZ, JL and ZG analyzed the computational and experimental results. JL wrote the manuscript under the supervision of LL and CZ.

## Acknowledgements

The authors gratefully acknowledge the National Center for Protein Sciences at Peking University in Beijing, China, for the platform of crystallization conditions screening and assistance of Dr. Tiantian Wei. The authors thank the high-performance computing platform of the Peking-Tsinghua Center for Life Sciences, Peking University, for providing the computational resources. The authors thank the National Facility for Protein Science in Shanghai (NFPS) and Shanghai Synchrotron Radiation Facility for the assistance of crystallographic data collection. The authors thank Jun Yang at Peking University in Beijing, China, for the assistance with X-ray diffraction data collection and analysis of 1FB0-3.

## References

[1] Sheng Wang, Siqi Sun, Zhen Li, Renyu Zhang, and Jinbo Xu. Accurate de novo prediction of protein contact map by ultra-deep learning model. PLoS Computational Biology, 13(1):e1005324, 2017.

[2] Andrew W Senior, Richard Evans, John Jumper, James Kirkpatrick, Laurent Sifre, Tim Green, Chongli Qin, Augustin Žídek, Alexander WR Nelson, Alex Bridgland, et al. Improved protein structure prediction using potentials from deep learning. Nature, 577(7792):706–710, 2020.

[3] Mirko Torrisi, Gianluca Pollastri, and Quan Le. Deep learning methods in protein structure prediction. Computational and Structural Biotechnology Journal, 18:1301–1310, 2020.

[4] Fandi Wu and Jinbo Xu. Deep template-based protein structure prediction. PLoS Computational Biology, 17(5):e1008954, 2021.

[5] Sergey Ovchinnikov, Hahnbeom Park, Neha Varghese, Po-Ssu Huang, Georgios A Pavlopoulos, David E Kim, Hetunandan Kamisetty, Nikos C Kyrpides, and David Baker. Protein structure determination using metagenome sequence data. Science, 355(6322):294–298, 2017.

[6] Gui-Jun Zhang, Lai-Fa Ma, Xiao-Qi Wang, and Xiao-Gen Zhou. Secondary structure and contact guided differential evolution for protein structure prediction. IEEE/ACM Transactions on Computational Biology and Bioinformatics, 17(3):1068–1081, 2018.

[7] Jianyi Yang, Ivan Anishchenko, Hahnbeom Park, Zhenling Peng, Sergey Ovchinnikov, and David Baker. Improved protein structure prediction using predicted interresidue orientations. Proceedings of the National Academy of Sciences, 117(3):1496–1503, 2020.

[8] Fusong Ju, Jianwei Zhu, Bin Shao, Lupeng Kong, Tie-Yan Liu, Wei-Mou Zheng, and Dongbo Bu. Copulanet: Learning residue co-evolution directly from multiple sequence alignment for protein structure prediction. Nature Communications, 12(1):1–9, 2021.

[9] Jinbo Xu, Matthew Mcpartlon, and Jin Li. Improved protein structure prediction by deep learning irrespective of co-evolution information. Nature Machine Intelligence, pages 1–9, 2021.

[10] John Jumper, Richard Evans, Alexander Pritzel, Tim Green, Michael Figurnov, Olaf Ronneberger, Kathryn Tunyasuvunakool, Russ Bates, Augustin Žídek, Anna Potapenko, et al. Highly accurate protein structure prediction with alphafold. Nature, 596(7873):583–589, 2021.

[11] Minkyung Baek, Frank DiMaio, Ivan Anishchenko, Justas Dauparas, Sergey Ovchinnikov, Gyu Rie Lee, Jue Wang, Qian Cong, Lisa N Kinch, R Dustin Schaeffer, et al. Accurate prediction of protein structures and interactions using a three-track neural network. Science, 373(6557):871–876, 2021.

[12] Brian Kuhlman and David Baker. Native protein sequences are close to optimal for their structures. Proceedings of the National Academy of Sciences, 97(19):10383–10388, 2000.

[13] Xiaoqiang Huang, Robin Pearce, and Yang Zhang. Evoef2: accurate and fast energy function for computational protein design. Bioinformatics, 36(4):1135–1142, 2020.

[14] Shide Liang, Zhixiu Li, Jian Zhan, and Yaoqi Zhou. De novo protein design by an energy function based on series expansion in distance and orientation dependence. Bioinformatics, 2021.

[15] Jingxue Wang, Huali Cao, John ZH Zhang, and Yifei Qi. Computational protein design with deep learning neural networks. Scientific Reports, 8(1):1–9, 2018.

[16] James O’Connell, Zhixiu Li, Jack Hanson, Rhys Heffernan, James Lyons, Kuldip Paliwal, Abdollah Dehzangi, Yuedong Yang, and Yaoqi Zhou. Spin2: Predicting sequence profiles from protein structures using deep neural networks. Proteins: Structure, Function, and Bioinformatics, 86(6):629–633, 2018.

[17] Sheng Chen, Zhe Sun, Lihua Lin, Zifeng Liu, Xun Liu, Yutian Chong, Yutong Lu, Huiying Zhao, and Yuedong Yang. To improve protein sequence profile prediction through image captioning on pairwise residue distance map. Journal of Chemical Information and Modeling, 60(1):391–399, 2019.

[18] John Ingraham, Vikas Garg, Regina Barzilay, and Tommi Jaakkola. Generative models for graph-based protein design. Advances in Neural Information Processing Systems, 32, 2019.

[19] Yuan Zhang, Yang Chen, Chenran Wang, Chun-Chao Lo, Xiuwen Liu, Wei Wu, and Jinfeng Zhang. Prodconn: Protein design using a convolutional neural network. Proteins: Structure, Function, and Bioinformatics, 88(7):819–829, 2020.

[20] Alexey Strokach, David Becerra, Carles Corbi-Verge, Albert Perez-Riba, and Philip M Kim. Fast and flexible protein design using deep graph neural networks. Cell Systems, 11(4):402–411, 2020.

[21] Bowen Jing, Stephan Eismann, Patricia Suriana, Raphael JL Townshend, and Ron Dror. Learning from protein structure with geometric vector perceptrons. arXiv preprint arXiv:2009.01411, 2020.

[22] Namrata Anand, Raphael Eguchi, Irimpan I Mathews, Carla P Perez, Alexander Derry, Russ B Altman, and Po-Ssu Huang. Protein sequence design with a learned potential. Nature Communications, 13(1):1–11, 2022.

[23] Yifei Qi and John ZH Zhang. Densecpd: improving the accuracy of neural-network-based computational protein sequence design with densenet. Journal of Chemical Information and Modeling, 60(3):1245–1252, 2020.

[24] Zhangyang Gao, Cheng Tan, and Stan Z Li. Pifold: Toward effective and efficient protein inverse folding. arXiv preprint arXiv:2209.12643, 2022.

[25] Milong Ren, Chungong Yu, Dongbo Bu, and Haicang Zhang. Highly accurate and robust protein sequence design with carbondesign. BioRxiv, pages 2023–08, 2023.

[26] Deniz Akpinaroglu, Kosuke Seki, Amy Guo, Eleanor Zhu, Mark JS Kelly, and Tanja Kortemme. Structure-conditioned masked language models for protein sequence design generalize beyond the native sequence space. BioRxiv, pages 2023–12, 2023.

[27] Yufeng Liu, Lu Zhang, Weilun Wang, Min Zhu, Chenchen Wang, Fudong Li, Jiahai Zhang, Houqiang Li, Quan Chen, and Haiyan Liu. Rotamer-free protein sequence design based on deep learning and self-consistency. Nature Computational Science, 2:451–462, 2022.

[28] Justas Dauparas, Ivan Anishchenko, Nathaniel Bennett, Hua Bai, Robert J Ragotte, Lukas F Milles, Basile IM Wicky, Alexis Courbet, Rob J de Haas, Neville Bethel, et al. Robust deep learning–based protein sequence design using proteinmpnn. Science, 378(6615):49–56, 2022.

[29] Nathaniel R Bennett, Brian Coventry, Inna Goreshnik, Buwei Huang, Aza Allen, Dionne Vafeados, Ying Po Peng, Justas Dauparas, Minkyung Baek, Lance Stewart, et al. Improving de novo protein binder design with deep learning. Nature Communications, 14(1):2625, 2023.

[30] Matthew McPartlon and Jinbo Xu. An end-to-end deep learning method for protein side-chain packing and inverse folding. Proceedings of the National Academy of Sciences, 120(23):e2216438120, 2023.

[31] Justas Dauparas, Gyu Rie Lee, Robert Pecoraro, Linna An, Ivan Anishchenko, Cameron Glasscock, and David Baker. Atomic context-conditioned protein sequence design using ligandmpnn. BioRxiv, pages 2023–12, 2023.

[32] Cyrus A Wilson, Julia Kreychman, and Mark Gerstein. Assessing annotation transfer for genomics: quantifying the relations between protein sequence, structure and function through traditional and probabilistic scores. Journal of Molecular Biology, 297(1):233–249, 2000.

[33] Annabel E Todd, Christine A Orengo, and Janet M Thornton. Evolution of function in protein superfamilies, from a structural perspective. Journal of Molecular Biology, 307(4):1113–1143, 2001.

[34] Ian Sillitoe, Nicola Bordin, Natalie Dawson, Vaishali P Waman, Paul Ashford, Harry M Scholes, Camilla SM Pang, Laurel Woodridge, Clemens Rauer, Neeladri Sen, et al. Cath: increased structural coverage of functional space. Nucleic Acids Research, 49(D1):D266–D273, 2021.

[35] Jiale Liu, Changsheng Zhang, and Luhua Lai. Geopacker: A novel deep learning framework for protein side-chain modeling. Protein Science, 31(12):e4484, 2022.

[36] Georgii G Krivov, Maxim V Shapovalov, and Roland L Dunbrack Jr. Improved prediction of protein side-chain conformations with scwrl4. Proteins: Structure, Function, and Bioinformatics, 77(4):778–795, 2009.

[37] Xiaoqiang Huang, Robin Pearce, and Yang Zhang. Faspr: an open-source tool for fast and accurate protein side-chain packing. Bioinformatics, 36(12):3758–3765, 2020.

[38] Magnus Ekeberg, Cecilia Lövkvist, Yueheng Lan, Martin Weigt, and Erik Aurell. Improved contact prediction in proteins: using pseudolikelihoods to infer potts models. Physical Review E, 87(1):012707, 2013.

[39] Sivaraman Balakrishnan, Hetunandan Kamisetty, Jaime G Carbonell, Su-In Lee, and Christopher James Langmead. Learning generative models for protein fold families. Proteins: Structure, Function, and Bioinformatics, 79(4):1061–1078, 2011.

[40] Patrick Conway, Michael D Tyka, Frank DiMaio, David E Konerding, and David Baker. Relaxation of backbone bond geometry improves protein energy landscape modeling. Protein Science, 23(1):47–55, 2014.

[41] Rebecca F Alford, Andrew Leaver-Fay, Jeliazko R Jeliazkov, Matthew J OMeara, Frank P DiMaio, Hahnbeom Park, Maxim V Shapovalov, P Douglas Renfrew, Vikram K Mulligan, Kalli Kappel, et al. The rosetta all-atom energy function for macromolecular modeling and design. Journal of Chemical Theory and Computation, 13(6):3031–3048, 2017.

[42] Wenkai Wang, Zhenling Peng, and Jianyi Yang. Single-sequence protein structure prediction using supervised transformer protein language models. Nature Computational Science, 2(12):804–814, 2022.

[43] Zeming Lin, Halil Akin, Roshan Rao, Brian Hie, Zhongkai Zhu, Wenting Lu, Nikita Smetanin, Robert Verkuil, Ori Kabeli, Yaniv Shmueli, et al. Evolutionary-scale prediction of atomic-level protein structure with a language model. Science, 379(6637):1123–1130, 2023.

[44] Ruidong Wu, Fan Ding, Rui Wang, Rui Shen, Xiwen Zhang, Shitong Luo, Chenpeng Su, Zuofan Wu, Qi Xie, Bonnie Berger, et al. High-resolution de novo structure prediction from primary sequence. BioRxiv, pages 2022–07, 2022.

[45] Domenico Cozzetto, Andriy Kryshtafovych, Michele Ceriani, and Anna Tramontano. Assessment of predictions in the model quality assessment category. Proteins: Structure, Function, and Bioinformatics, 69(S8):175–183, 2007.

[46] Björn Wallner. AFsample: improving multimer prediction with AlphaFold using massive sampling. Bioinformatics, 39(9):btad573, 09 2023.

[47] Limin Fu, Beifang Niu, Zhengwei Zhu, Sitao Wu, and Weizhong Li. Cd-hit: accelerated for clustering the next-generation sequencing data. Bioinformatics, 28(23):3150–3152, 2012.

[48] Naozumi Hiranuma, Hahnbeom Park, Minkyung Baek, Ivan Anishchenko, Justas Dauparas, and David Baker. Improved protein structure refinement guided by deep learning based accuracy estimation. Nature Communications, 12(1):1340, 2021.

[49] Xavier Robin, Juergen Haas, Rafal Gumienny, Anna Smolinski, Gerardo Tauriello, and Torsten Schwede. Continuous automated model evaluation (cameo)perspectives on the future of fully automated evaluation of structure prediction methods. Proteins: Structure, Function, and Bioinformatics, 89(12):1977–1986, 2021.

[50] Bin Huang, Tingwen Fan, Kaiyue Wang, Haicang Zhang, Chungong Yu, Shuyu Nie, Yangshuo Qi, Wei-Mou Zheng, Jian Han, Zheng Fan, et al. Accurate and efficient protein sequence design through learning concise local environment of residues. Bioinformatics, 39(3):btad122, 2023.

[51] Ivan Anishchenko, Samuel J Pellock, Tamuka M Chidyausiku, Theresa A Ramelot, Sergey Ovchinnikov, Jingzhou Hao, Khushboo Bafna, Christoffer Norn, Alex Kang, Asim K Bera, et al. De novo protein design by deep network hallucination. Nature, 600(7889):547–552, 2021.

[52] Joseph L Watson, David Juergens, Nathaniel R Bennett, Brian L Trippe, Jason Yim, Helen E Eisenach, Woody Ahern, Andrew J Borst, Robert J Ragotte, Lukas F Milles, et al. De novo design of protein structure and function with rfdiffusion. Nature, 620(7976):1089–1100, 2023.

[53] John B Ingraham, Max Baranov, Zak Costello, Karl W Barber, Wujie Wang, Ahmed Ismail, Vincent Frappier, Dana M Lord, Christopher Ng-Thow-Hing, Erik R Van Vlack, et al. Illuminating protein space with a programmable generative model. Nature, pages 1–9, 2023.

[54] Alexander E Chu, Lucy Cheng, Gina El Nesr, Minkai Xu, and Po-Ssu Huang. An all-atom protein generative model. BioRxiv, pages 2023–05, 2023.

[55] Guido Capitani, Zora Marković-Housley, Gregoire DelVal, May Morris, Johan N Jansonius, et al. Crystal structures of two functionally different thioredoxins in spinach chloroplasts. Journal of Molecular Biology, 302(1):135–154, 2000.

[56] Zeming Lin, Halil Akin, Roshan Rao, Brian Hie, Zhongkai Zhu, Wenting Lu, Nikita Smetanin, Robert Verkuil, Ori Kabeli, Yaniv Shmueli, Allan dos Santos Costa, Maryam Fazel-Zarandi, Tom Sercu, Salvatore Candido, and Alexander Rives. Evolutionary-scale prediction of atomic-level protein structure with a language model. Science, 379(6637):1123–1130, 2023.

